# Time-course analysis of *Streptococcus sanguinis* after manganese depletion reveals changes in glycolytic, nucleotide, and redox metabolites

**DOI:** 10.1101/2020.08.30.274233

**Authors:** Tanya Puccio, Biswapriya B. Misra, Todd Kitten

**Author notes:** Corresponding author: Todd Kitten.

## Abstract

**Introduction:** Manganese is important for the endocarditis pathogen, *Streptococcus sanguinis*. Little is known about why manganese is required for virulence or how it impacts the metabolome of streptococci.

**Objectives:** We applied untargeted metabolomics to cells and media to understand temporal changes resulting from manganese depletion.

**Methods:** EDTA was added to a *S. sanguinis* manganese-transporter mutant in aerobic fermentor conditions. Cell and media samples were collected pre- and post-EDTA treatment. Metabolomics data were generated using positive and negative modes of data acquisition on an LC-MS/MS system. Data were subjected to statistical processing using MetaboAnalyst and time-course analysis using Short Time series Expression Miner (STEM).

**Results:** We observed quantitative changes in 534 and 422 metabolites in cells and media, respectively, after EDTA addition. The 173 cellular metabolites identified as significantly different indicated enrichment of purine and pyrimidine metabolism. Further multivariate analysis revealed that the top 15 cellular metabolites belonged primarily to lipids and redox metabolites. The STEM analysis revealed global changes in cells and media in comparable metabolic pathways. Products of glycolysis such as pyruvate and fructose-1,6-bisphosphate increased, suggesting that enzymes that act on them may require manganese for activity or expression. Nucleosides accumulated, possibly due to a blockage in conversion to nucleobases. Simultaneous accumulation of *ortho*-tyrosine and reduced glutathione suggests that cells were unable to utilize glutathione as a reductant.

**Conclusion:** Differential analysis of metabolites revealed the activation of a number of metabolic pathways in response to manganese depletion, many of which may be connected to carbon catabolite repression.

## 1. Introduction

*Streptococcus sanguinis* is a gram-positive bacterium known for its duplicity. As an early and abundant colonizer of teeth, *S. sanguinis* is associated with oral health (Kreth *et al*., 2017; Kreth *et al*., 2005). However, when it enters the bloodstream, whether through dental procedures or activities as routine as eating, it is known to colonize the heart valves or other endocardial surfaces of persons with particular pre-existing cardiac conditions, leading to infective endocarditis (IE) (Moreillon *et al*., 2002; Widmer *et al*., 2006). IE has a global mortality rate of 12-40% (Bor *et al*., 2013; Cahill *et al*., 2017). Historically, prevention has relied upon administration of prophylactic broad-spectrum antibiotics to high-risk patients prior to dental visits (Wilson *et al*., 2007). With rising antibiotic resistance (Dodds, 2017), as well as controversial efficacy (Quan *et al*., 2020; Thornhill *et al*., 2018), novel drug targets that are required for endocarditis causation but not beneficial colonization are under investigation.

One such potential drug target in *S. sanguinis* is the lipoprotein SsaB, a component of the ATP-binding cassette transporter SsaACB. This transporter and orthologs in related species have been shown to be important for manganese (Mn) transport and virulence (Colomer-Winter *et al*., 2018; Crump *et al*., 2014; Dintilhac *et al*., 1997; Kehl-Fie *et al*., 2013). Previous studies utilizing a Δ*ssaACB* strain of *S. sanguinis* revealed that this mutant is deficient in cellular Mn levels (Murgas *et al*., 2020) and virulence in our rabbit model of IE (Baker *et al*., 2019). These studies also suggested that the reduced virulence of Mn-deficient cells is due to growth arrest in the aerobic, low-Mn environment characteristic of an aortic valve infection, implying the existence of one or more Mn-dependent metabolic pathways that are essential for aerobic growth. The metabolic pathway(s) and individual metabolites involved have not been defined.

Metabolomics is the comprehensive study of small molecules in the molecular weight range of 50-2000 Da in biological systems. Diverse mass spectrometry platforms such as LC-MS/MS, GC-MS and CE-MS with and without chromatography, and spectroscopy technologies such as NMR have enabled high-throughput discovery metabolomics in various biological systems, including bacteria, plants, and humans (Misra and Olivier, 2020). Recent studies have described the metabolomes of certain streptococci using various mass spectrometry methods: *Streptococcus intermedius* under various oxygen conditions (Fei *et al*., 2016); *Streptococcus pneumoniae* in chemically defined medium (Leonard *et al*., 2018); and *Streptococcus thermophilus* in pH-controlled batch fermentation (Liu *et al*., 2020; Qiao *et al*., 2019). To our knowledge, the metabolome of *S. sanguinis* has yet to be investigated. Here we report the first untargeted metabolomic analysis of *S. sanguinis* or, indeed, of any *Streptococcus*, under Mn replete vs. deplete conditions.

## 2. Materials and Methods

### 2.1 Bacterial strains and growth conditions

*S. sanguinis* strain SK36 was isolated from human dental plaque (Kilian *et al*., 1989; Xu *et al*., 2007). The Δ*ssaACB* strain (JFP169) was generated from SK36 previously by replacement of the *ssaACB* genes with the *aphA-3* gene encoding kanamycin resistance (Puccio *et al*., 2020). Overnight pre-cultures were created by inoculation of Brain Heart Infusion (BHI) broth (Becton, Dickinson and Company, Franklin Lakes, NJ) with single-use aliquots of cryopreserved cells by 1000-fold dilution. Kanamycin (Sigma-Aldrich, St. Louis, MO) was added to 500 µg mL^−1^ for Δ*ssaACB* pre-cultures. Pre-cultures were incubated at 37°C for 18 h in 6% O_2_ (6% O_2_, 7% H_2_, 7% CO_2_ and 80% N_2_) using an Anoxomat (Advanced Instruments, Norwood, MA) jar.

### 2.2 Fermentor growth conditions and sample collection

Aerobic fermentor growth of Δ*ssaACB* cell culture was achieved using a BIOSTAT® B bioreactor (Sartorius Stedim, Göttingen, Germany) and samples were collected as described in Puccio and Kitten (2020). The pre-EDTA sample was collected at T_-20_ (min), where EDTA was added at T_0_ to a final concentration of 100 µM. Post-EDTA samples were collected at T_25_ and T_50_. All samples were stored at −80°C until shipped on dry ice to Metabolon, Inc. (Durham, North Carolina) for further analysis.

### 2.3 Sample preparation, UPLC-MS/MS, data extraction, compound identification, and curation

Metabolomics sample processing was completed by Metabolon, Inc. as described in the Supplementary Methods and in previous publications (Dehaven *et al*., 2010; Evans *et al*., 2009).

### 2.4 Statistical analysis of metabolomics and transcriptomics datasets

Statistical analysis of the metabolomics data sets was performed using statistical software R (Version 3.5.2) (Team, 2018). Normalized, transformed, imputed, outlier-removed, and scaled-peak areas representative of relative metabolite amounts obtained using DeviumWeb (Grapov, 2014) are presented. Hierarchical clustering analysis (HCA) was performed on Pearson distances using MetaboAnalyst 4.0 (www.metaboanalyst.ca) (Xia *et al*., 2015), with the data normalized using z-scores of the relative abundance of the metabolites for heat map display. Correlations reported are Spearman rank correlations. Principal component analysis (PCA) and partial least squared discriminant analyses (PLS-DA) were performed using MetaboAnalyst, with the output displayed as score plots for visualization of sample groups. One-way analysis of variance (ANOVA) followed by post-hoc analysis using Fisher’s least significant difference (LSD) test was used for analysis of statistical significance using MetaboAnalyst.

### 2.5 Time-course analysis of cellular and media metabolomes

For our 70 min time course, we used the Short Time series Expression Miner (STEM) tool. The following parameters were used for our analysis: no normalization of data; 0 added as the starting point; number of model profiles = 20; maximum unit change in model profiles between time points = To explain the model profiles, we used an expression of −1 if levels of a metabolite decreased, 0 if levels were unchanged, and 1 if levels increased. For instance, a model profile with an expression of −1, −1, 0, represents decreased, decreased, and unchanged, levels of a given set of metabolites for the 3 time points.

### 2.6 Metabolic pathway and enrichment analysis

Pathway enrichment analysis was performed using MetaboAnalyst 4.0 and reported pathways are KEGG-based (Kanehisa and Goto, 2000). The Chemical Translation Service (CTS: http://cts.fiehnlab.ucdavis.edu/conversion/batch) was used to convert the common chemical names into their KEGG, HMDB, Metlin, PubChem CID, and ChEBI identifiers.

## 3. Results

### 3.1 EDTA treatment of Δ*ssaACB* cells leads to Mn depletion and slowed growth

As described in Puccio *et al*. (2020), EDTA treatment of Δ*ssaACB* aerobic fermentor-grown cells results in the depletion of Mn but no other biologically relevant metals, such as Fe or Zn, as determined by inductively coupled plasma optical emission spectroscopy (ICP-OES) **(Figure 1)**. Beginning ∼38 min post-EDTA addition, cell growth slowed, resulting in a steady drop in OD **(Figure 1)**.

**Fig. 1.**
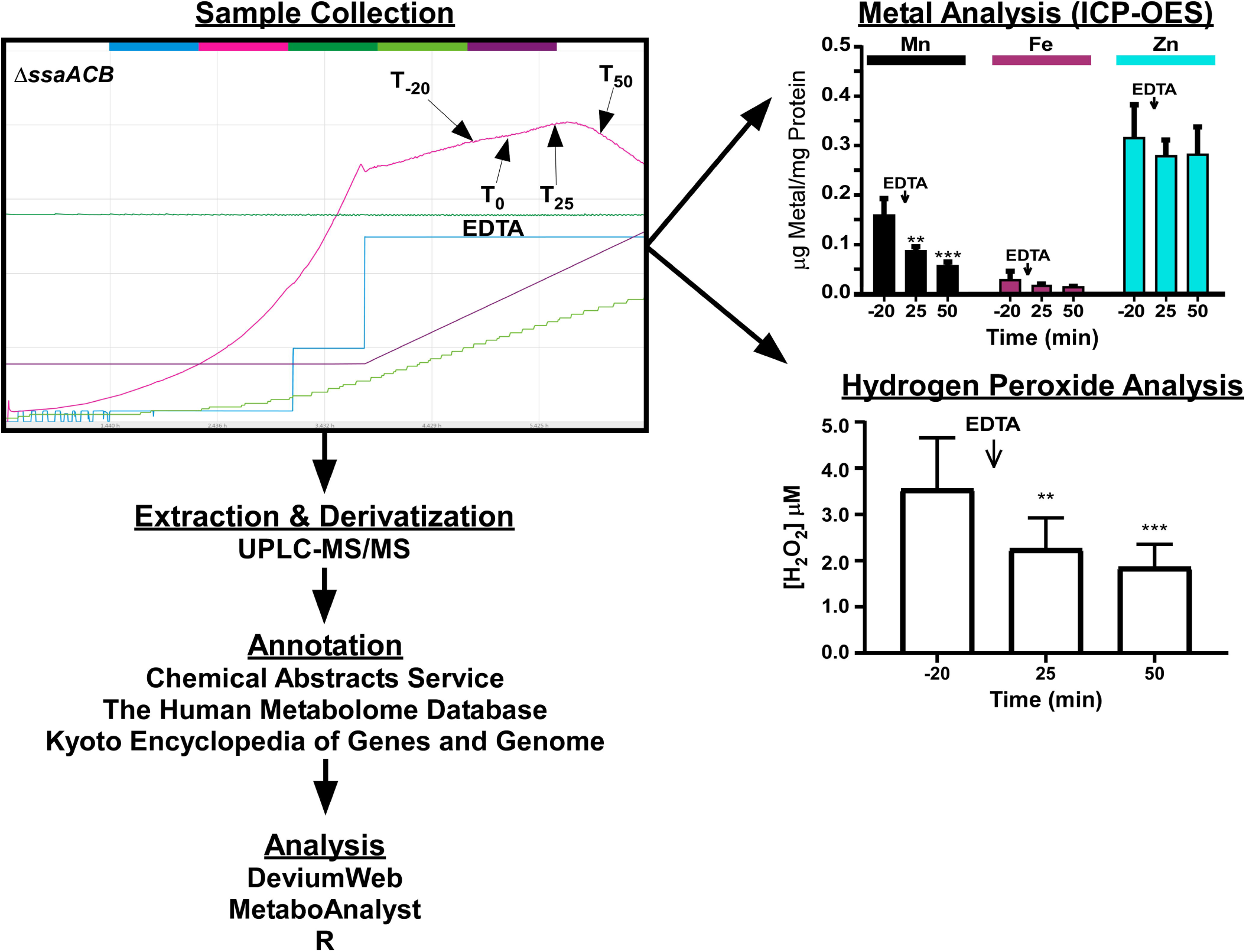
Schematic diagram displaying the experimental design, platform and software tools used for the analysis of metabolomic changes in cells and media subjected to EDTA treatment. Fermentor sample collection, metal, and hydrogen peroxide analysis charts were adapted from Puccio, et al. (2020). Extraction, derivatization, and annotation were completed by Metabolon, Inc. ICP-OES, inductively coupled plasma optical emission spectroscopy; UPLC-MS/MS, ultra performance liquid chromatography with tandem mass spectrometry.

### 3.2 Global metabolomics of *S. sanguinis* cells and BHI media

Our goal was to understand the metabolic consequences of Mn depletion during growth of a *S. sanguinis* Mn-transporter mutant in a rich medium (BHI), as well as to survey changes in the conditioned media during the growth and treatment periods. Extensive global untargeted metabolomics analysis revealed 534 metabolites in cells and 422 metabolites in conditioned media. The raw metabolite abundance values alongside the identified metabolite IDs, super pathways and sub-pathway names, average mass, and identifiers such as Chemical Abstracts Service (CAS), PubChem, Kyoto Encyclopedia of Genes and Genomes (KEGG), and Human Metabolome Database (HMDB) IDs are provided for both cellular and media metabolites (**Tables S1-S2**). These datasets were refined through normalization, transformation, and scaling, followed by imputation (**Tables S3-S4)**. The 534 metabolites belong to 57 different KEGG metabolic pathways (**Table S5**). The 422 metabolites quantified in the conditioned BHI media belonged to 50 different metabolic pathways (**Table S6**), all of which overlap with the metabolic pathways found in the cells.

BHI has as its chief constituents bovine and porcine brain and heart extracts. Based on comparison with the pre-inoculation media samples, we identified several metabolites that appear to originate from BHI, and were excluded from further statistical processing as they were unique to the growth media alone **(Table S7)**. Any metabolite that occurred in fewer than 75% of the samples was also excluded from the analysis, which resulted in the exclusion of 9 of the 534 metabolites detected in cells **(Table S7)**.

### 3.3 Differential accumulation patterns of metabolites over time course and EDTA treatment

We used a false discovery rate (FDR)-corrected ANOVA to determine metabolites that were significantly different in abundance between the different time-points. ANOVA revealed 173 and 13 metabolites that were significantly different in cells and media, respectively **(Tables S8-9)**. To investigate whether these differential metabolites would map to metabolic pathways, we mapped the set of metabolites using the *Streptococcus pyogenes* M1 476 KEGG database within MetaboAnalyst by implementing overrepresentation analysis with Fisher’s exact test and pathway topology analysis using relative-betweenness centrality (Jewison *et al*., 2014). Pathway enrichment analysis of the 173 cellular metabolites that were differential along the time course of EDTA treatment identified only purine and pyrimidine metabolism (nominal P-value < 0.05) (**Figure S1a; Table S10)**. Surprisingly pathway enrichment analysis of the 13 media metabolites that were differential along the time course identified purine and pyrimidine metabolism as above, but also glyoxylate and dicarboxylate metabolism, and alanine, aspartate, and glutamate metabolism (nominal P-value < 0.05) (**Figure S1b; Table S11**). When metabolite abundances were compared for the two post-EDTA time points vs T_-20_, it was revealed that one, five, 13, and 30 metabolites were increased in T_25_ and T_50_ in media and T_25_ and T_50_ in cells, respectively **(Figure S1c)**. Of these, only 2’-deoxyadenosine increased in both cells and media at T_50_ (**Tables S12-13**). The 30 metabolites increased in T_50_ in cells were mostly lipids, energy metabolites, nucleotide phosphates, and dinucleotides (**Table S12**). When significantly decreased metabolites were compared, it was revealed that 1, 1, 13, and 30 metabolites were decreased in T_25_ and T_50_ in media and T_25_ and T_50_ in cells, respectively (**Figure S1d**). Only glutamine levels decreased in both media samples (**Table S13**). The five metabolites that decreased in cells at T_25_ included cCMP and cUMP, while the 18 metabolites that decreased at T_50_ in the cells included IMP and XMP (**Table S12**).

### 3.4 Multivariate and hierarchical clustering analysis

To define the metabolomic changes caused by Mn depletion, we used multivariate analysis and HCA. Using an unsupervised multivariate analysis, PCA, we observed that metabolite abundances alone were able to discriminate between the samples and explain 58.8% of the variation in the dataset by virtue of the first 2 PCs (PC1, PC2) in cells (**Figure 2a)** and 67.5% in media **(Figure 2b)**.

**Fig. 2.**
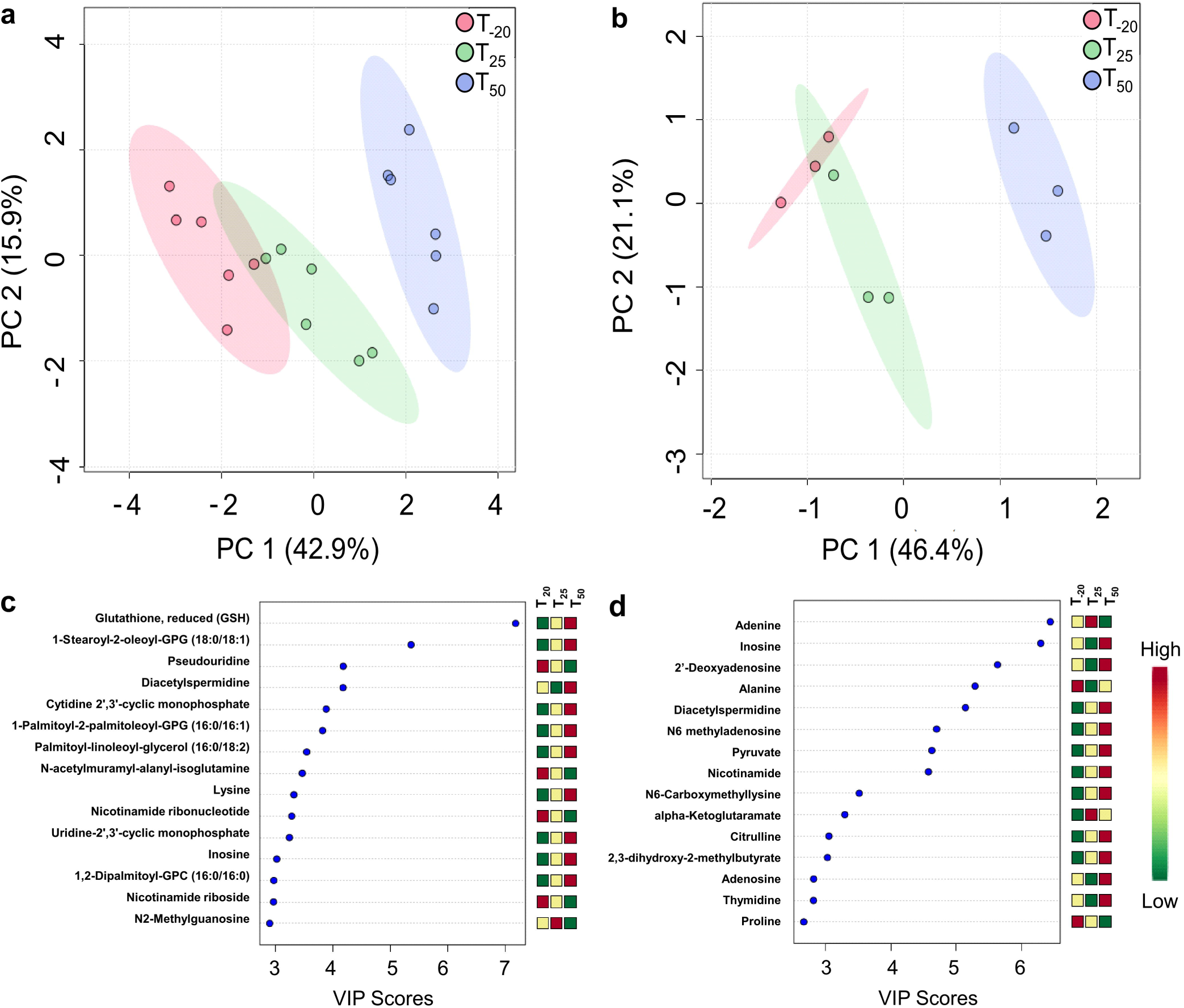
Multivariate, VIP, and time course analysis of the metabolomic changes in cells and media. Score plots of PCA displaying the separation of time-points in cells (a) and spent media (b). Cell samples n = 6; media samples n = 3. Top 15 metabolites (variables) based on VIP scores from PLS-DA analysis of cells (c) and spent media (d).

Using supervised multivariate analysis, PLS-DA, we observed that metabolite abundances alone were able to discriminate between the samples and explain 57.1 % of the variation in the dataset by virtue of the first 2 PCs (Component 1 and 2) in cells (**Figure S2a)** and 57.7% in media **(Figure S2b)**. Additionally, PLS-DA and PCA performed on spent media samples explained 93.4% and 93.5% of the variation, respectively, by virtue of the first 2 PCs **(Figure S2c-d)**.

To identify the metabolites responsible for the discrimination among the metabolomic profiles, the variable importance in projection (VIP) score was used to select features with the most significant contribution in a PLS-DA model. VIP scores are a weighted sum of PLS weights for each variable and measure the contribution of each predictor variable to the model. Further, the VIP statistic summarizes the importance of the metabolites in differentiating the sample time points in multivariate space. Metabolites exhibiting high VIP scores (≥1.5) are the more influential variables. Our VIP analysis revealed that the top 15 metabolites for cells included lipids, cCMP, cUMP, and redox metabolites **(Figure 2c)**. The VIP analysis revealed that the top 15 metabolites for spent media included amino acids and organic acids **(Figure 2d)**. Of these VIP metabolites (cut off ≥1.5), seven (glutamine, adenosine, adenine, glycerate, forminoglutamate, citrulline, and orotate) were shared between cells and media across all the time points, indicating their importance.

We performed an HCA using the z-score-normalized metabolite abundances of the cellular and media metabolites, separately **(Figure S3)**. Results indicated a clear clustering for the three time points as shown for the top 25 metabolites obtained from the ANOVA for individual sample groups. In cells, two distinct clusters were formed based on the metabolite abundances, where the upper cluster (decreased in T_50_) was represented by acetylated metabolites, purines and pyrimidines, and glutamyl dipeptides, and the bottom cluster (increased in T_50_) contained several amino acids and lipids, and cCMP, cUMP, and UTP **(Figure S3a)**. In media, two distinct clusters were formed based on the metabolite abundances, with the upper cluster (increased in T_50_) represented by several important metabolites such as uracil, ribose, pyruvate, nicotinamide, inosine, adenosine, guanosine, and the bottom cluster (decreased in T_50_) containing glutamine, adenine, and 3’AMP **(S3b)**.

### 3.5 Time-course analysis of cellular and media metabolites

To understand the time course-dependent changes in metabolite accumulation patterns across the three time points in this complex study design, we started with a clustering analysis. Using STEM analysis, we interrogated the time course changes of the metabolites in the cells and media. The metabolite abundances were put into 20 model clusters, which revealed differential accumulation of metabolites as a function of time. For the cells, the top two significant models were #19 (pattern 0, 1, 1, −1, P-value 5e-115) and #18 (pattern 0, 1, −1, 0, P-value 4e-12) representing 193 and 80 metabolites, respectively **(Figure S4a; Table S14)**. Metabolites following the pattern in model #19 were enriched for amino acid metabolic pathways: valine, leucine and isoleucine biosynthesis and degradation, alanine, aspartate and glutamate metabolism, and glycine, serine and threonine metabolism (P-value, < 0.1). Model #18 metabolites were enriched for arginine biosynthesis, arginine and proline metabolism, histidine metabolism, glyoxylate and dicarboxylate metabolism, and pyrimidine metabolism (P-value < 0.1). For the media, the top three models were #18 (0, 1, −1, 0, P-val-3e-59), #19 (pattern 0, 1, 1, −1, P-val-3e-23) and #14 (pattern 1, 1, 1, 1, P-val-6e-24) representing 132, 81, and 4 metabolites, respectively **(Figure S4b and Table S15)**. Metabolites following the pattern in model #18 were enriched for alanine, aspartate and glutamate metabolism, amino acid metabolism, and arginine and proline metabolism. Those in model #19 were enriched for arginine biosynthesis, valine, leucine and isoleucine biosynthesis and degradation, glyoxylate and dicarboxylate metabolism, pyrimidine metabolism, alanine, aspartate and glutamate metabolism, and glycine, serine and threonine metabolism. The metabolites in model #14 included 2-deoxyadenosine, N6-methyladenosine, inosine, and nicotinamide.

## 4. Discussion

### 4.1 Metabolomic analysis of BHI spent media reveals metabolic interactions of *S. sanguinis* with the extracellular environment

Our purpose in conducting this study was to examine the role of Mn in *S. sanguinis* metabolism, particularly in relation to IE. While the perfect medium for such a study would have been serum or plasma, this would not have been feasible, and so we instead used another complex yet commercially accessible medium—BHI. As with plasma, BHI has glucose as its most abundant sugar (0.2% w/v in BHI and ∼0.1% w/v in plasma). Although serum and plasma have been the subject of many metabolomic studies, we are not aware of any previous metabolomic analysis of BHI. Thus, the analysis of the pre-inoculated BHI (**Table S2**) may be of interest to the many investigators who use this medium. Likewise, the comparison of the pre-inoculated and T_-20_ media samples tells us much concerning the metabolic and transport capabilities of *S. sanguinis* under Mn-replete conditions (**Table S13**).

As expected, we observed a significant decrease of glucose in spent media (**Figure 3a**), indicating its utilization as carbon source. Levels of fructose and mannose significantly decreased as well **(Figure 3a)**, indicating that they are catabolized by cells. *S. sanguinis* encodes a number of putative sugar transport systems (Ajdic and Pham, 2007; Xu *et al*., 2007). Lactate and pyruvate levels increased significantly in the media after cell growth (**Figure 3b**), indicating that these products of glycolysis were secreted from cells. Pyruvate has been shown to be secreted by *S. sanguinis*, presumably to protect the cells from H_2_O_2_ stress by acting as an antioxidant (Redanz *et al*., 2020).

**Fig. 3.**
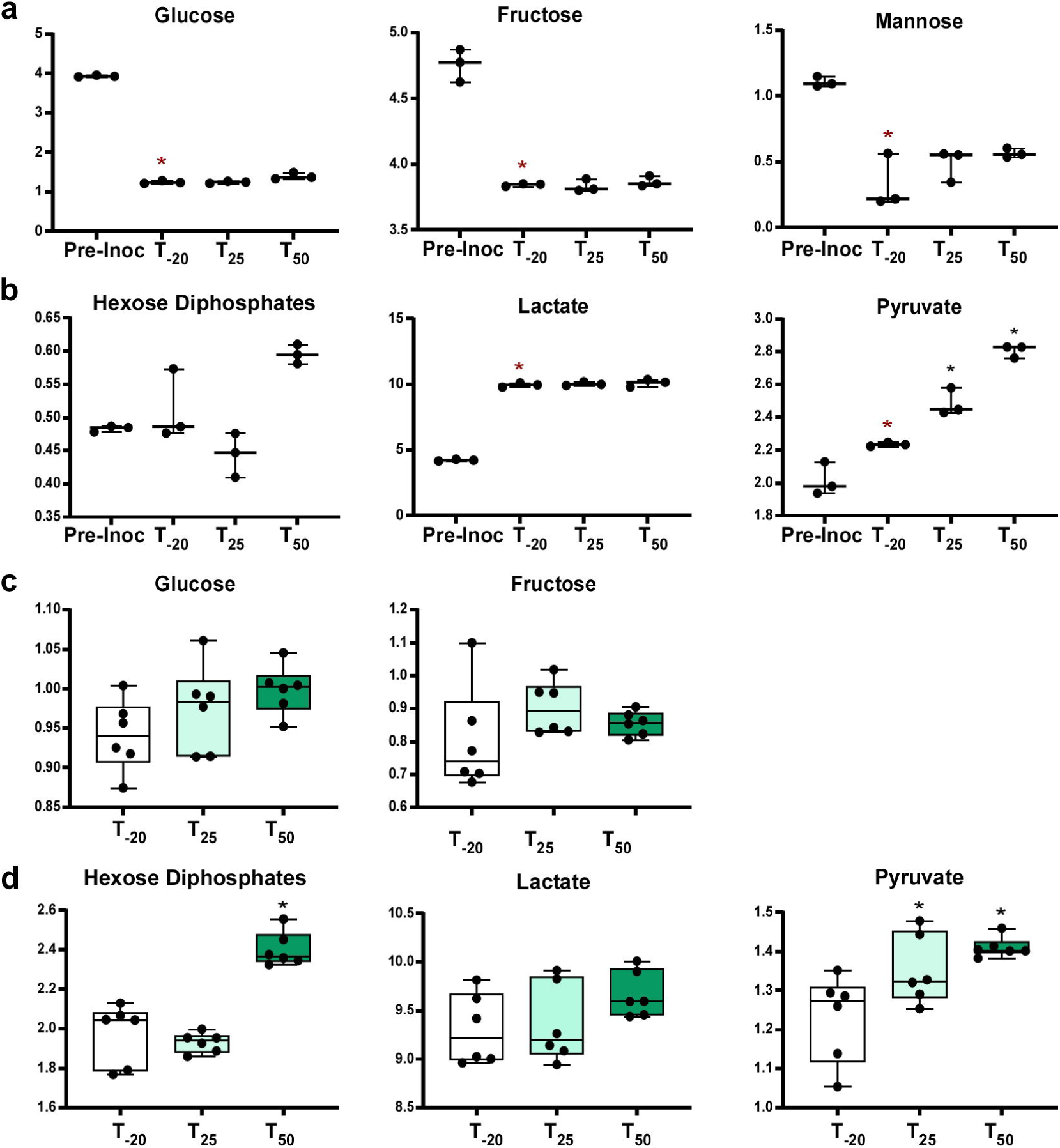
Relative abundance of carbohydrates and glycolytic intermediates in media and cells. Levels of sugars in media (a) and cells (c) are depicted. Products of glycolysis in media (b) and cells (d). Whiskers indicate the range; horizontal bars represent the mean. A two-tailed t-test was used to compare the pre-inoculum (Pre-Inoc) media samples to post-inoculum (T_-20_). Red asterisks indicate P-value < 0.05. Spent media and cell metabolite levels were compared using one-way ANOVA with a Fisher’s least significant difference test to compare the post-EDTA samples to pre-EDTA. Black asterisks indicate P-value < 0.05.

Also of interest, all nucleosides were significantly decreased after *S. sanguinis* growth (**Figures 4 and S5a-b**). The opposite trend was observed with nucleobases, where most were significantly increased after cell growth (**Figures 4 and S5c-d**). Nucleoside transport for salvage has been characterized in many bacteria, including the related species *Lactococcus lactis* (Martinussen *et al*., 2010) and *Streptococcus mutans* (Webb and Hosie, 2006).

**Fig. 4.**
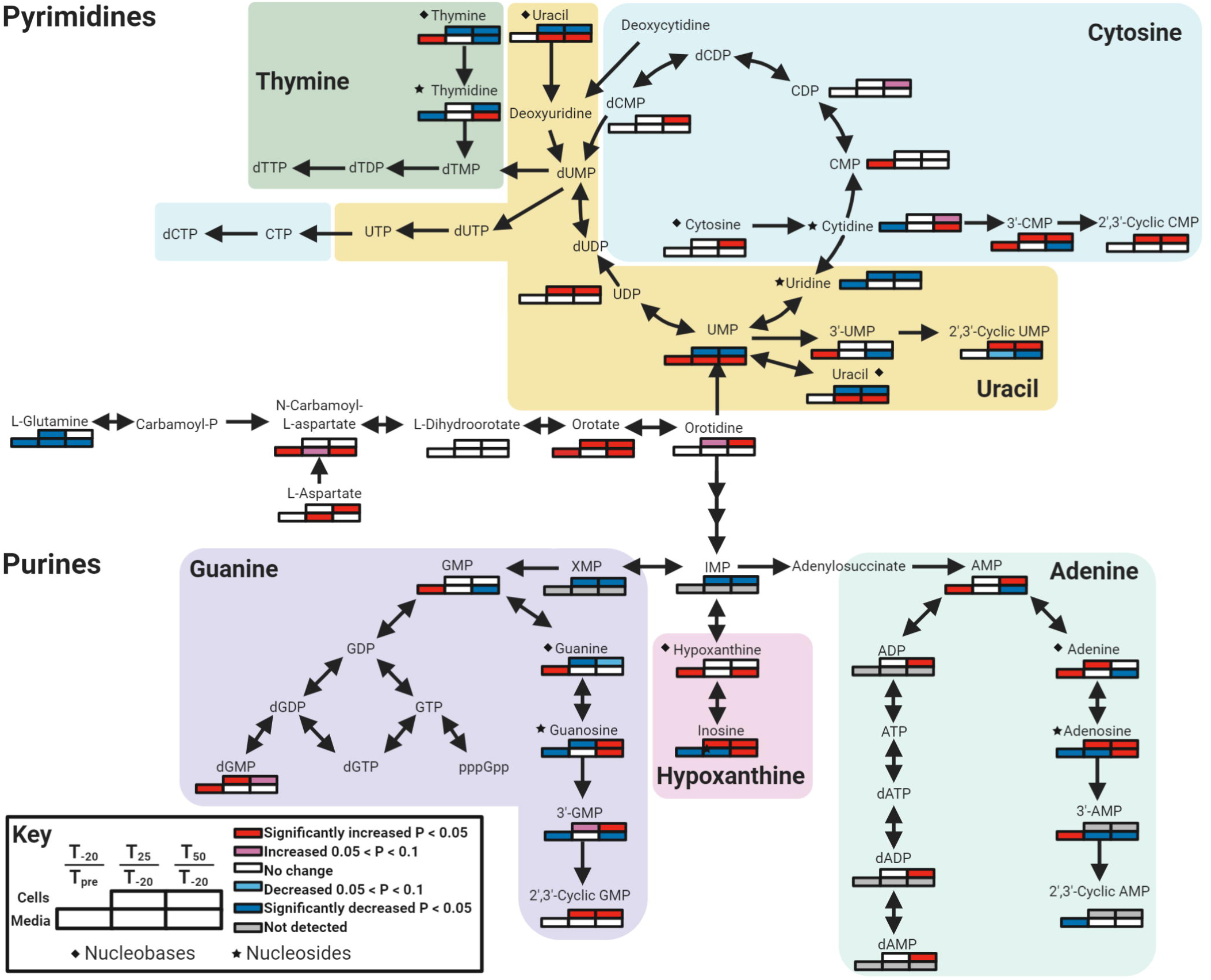
Quantitative changes in nucleotide metabolism for cells and media after Mn depletion. The direction of change in metabolite concentration is depicted in shades of red or blue, for increasing or decreasing concentration, respectively. Significance was determined by a t-test using the comparisons shown in the key. Metabolites that do not have a set of boxes were not detected in any sample. Diamonds indicate nucleobases and stars indicate nucleosides. Figure was made using Biorender.com.

### 4.2 Carbohydrate metabolism and glycolytic regulation in *S. sanguinis* cells show Mn dependence

The levels of glycolytic byproducts in *S. sanguinis* cells and spent media were impacted by Mn depletion. Glucose, fructose, and lactate levels remained constant in cells at all three time points while pyruvate levels increased after Mn depletion **(Figure 3c-d)**. Mannose was not detected in cells at any time point, indicating rapid catabolism by cells (**Table S1**). Lactate is known to be produced in high levels by streptococci and other lactic acid bacteria (Jakubovics *et al*., 2014), which explains the observed increase of lactate in the media after cellular growth. Pyruvate is produced through metabolism of sugars or amino acids. The observed increase in pyruvate levels in cells after Mn depletion **(Figure 4b)** is not due to increased sugar levels, as the flow of media remained constant throughout the experiment. Most amino acid levels remained unchanged or decreased in cells after Mn depletion (**Table S11**). One potential explanation for the increase in pyruvate levels is that fewer pyruvate molecules were oxidized by pyruvate oxidase (SpxB) into H_2_O_2_ and acetyl phosphate, consistent with our finding of a significant decrease in H_2_O_2_ levels after Mn depletion (**Figure 1**) (Puccio *et al*., 2020).

There was a significant accumulation of hexose diphosphates in cells at T_50_ and a slight increase in spent media as well (**Figure 3b & d**). Since levels of other glycolytic intermediates such as glucose-6-phosphate, glycerone, and glyceraldehyde-3-phosphate could not be measured using our platform **(Tables S1-2)**, we are unable to assess the impact on this pathway using metabolomics alone. We hypothesize that the hexose diphosphate is primarily fructose-1,6-bisphosphate and its accumulation results from the reduced activity of two potentially Mn-cofactored fructose-1,6-bisphosphate-consuming enzymes in the glycolytic pathway: fructose-1,6-bisphosphatase (Fbp; SSA_1056) and fructose-bisphosphate aldolase (Fba; SSA_1992) (Puccio *et al*., 2020). We further hypothesize that fructose-1,6-bisphosphate accumulation is at least partly responsible for the glucose-independent CcpA repression observed in the transcriptome of *S. sanguinis* after Mn depletion (Puccio *et al*., 2020).

Previous studies with other bacteria support a role for Mn in carbon metabolism. Mn deprivation was previously found to induce flux to the pentose phosphate pathway in *S. pneumoniae* (Ogunniyi *et al*., 2010). *Staphylococcus aureus* was found to be more susceptible to calprotectin-mediated Mn starvation when glucose was the sole carbon source than when amino acids were also present (Radin *et al*., 2016). Excess Mn modulated glycolysis in *Escherichia coli* biofilms by decreasing levels of glucose-6-phosphate and glyceraldehyde-3-phosphate (Guo and Lu, 2020). Here we provide further evidence that Mn levels impact central carbon metabolism.

### 4.3 Purine and pyrimidine metabolism in Mn-deplete *S. sanguinis* reveal nucleoside utilization from media and nucleobase accumulation in cells

Mn is known to impact nucleotide metabolism through its role as cofactor for the aerobic ribonucleotide reductase NrdF (Makhlynets *et al*., 2014; Rhodes *et al*., 2014). Here, we observed further impacts of Mn on nucleotide metabolism. Mean levels of guanosine, inosine, and adenosine increased in both cells and media at T_50_ (**Figures 4 and S5a & e**). In cells, guanine levels decreased while hypoxanthine and adenine levels were unchanged at T_50_ **(Figures 4 and S5g**). This indicates that there may be blockages in the conversion of purine nucleobases into nucleosides. There are three enzymes encoded by *S. sanguinis* that can catalyze this reaction: PunA (SSA_1258), DeoD (SSA_1259), and SSA_2046. None of these enzymes have been found to use Mn (BRENDA https://www.brenda-enzymes.org/) (Jeske *et al*., 2019). In our recent transcriptomics study, expression of *punA* and *deoD* were significantly decreased after Mn depletion (Puccio *et al*., 2020). The operon encoding *deoD* and *punA* has a carbon responsive element (*cre*) upstream of the first gene, *rpiA* (Bai *et al*., 2019), which is the recognition sequence for the carbon catabolite repression (CCR) regulator CcpA (Warner and Lolkema, 2003). As observed in Puccio *et al*. (2020), Mn depletion results in many changes in the CcpA regulon, which may explain the repression of this operon at T_50_. Thus, this may be but one example of a non-carbon catabolite pathway impacted by Mn depletion through its effect on CCR.

Similar to the purines, the pyrimidine nucleosides appear to be taken up from the media and the nucleobases were likely generated by cells **(Figures 4 and S5)**. Mean uridine levels in cells decreased slightly in cells after Mn depletion, whereas UMP **(Figure 4**) and uracil (**Figures 4 and S5h**) levels dropped significantly. Uracil levels in cells likely decreased due to lower UMP production. Interestingly, orotidine levels increased in cells (**Figure 4**), indicating a potential blockage in the conversion to UMP, although the explanation for this remains elusive as no PyrF enzyme is known to utilize a Mn cofactor (https://www.brenda-enzymes.org/).

Levels of thymine decreased in cells after Mn depletion (**Figures 4 and S5d & h**). which corresponds to a decrease in expression of *pdp* (pyrimidine nucleoside phosphorylase; SSA_1035; thymidine to thymine conversion) (Puccio *et al*., 2020). Oddly, thymidine levels decreased as well, although this may be explained by the increase in dTDP-rhamnose levels at T_50_ (**Table S10**), indicating that thymidine may have been shuttled to sugar metabolism. Mean cytosine and cytidine levels increased slightly in cells after Mn depletion **(Figures 4 and S5f & h**), which is the opposite trend from the other pyrimidines. Levels of downstream products 3’-CMP and 2’, 3’-cyclic CMP increased as well (**Figure 4**). The discrepancy may be explained by decreased conversion to uridine as its levels dropped after Mn depletion **(Figures 4 and S5f**). This is supported by a decrease in expression of *cdd* (cytidine deaminase; SSA_1037) after Mn depletion (Puccio *et al*., 2020) and Cdd may be Mn-cofactored (Hosono and Kuno, 1973).

### 4.4 Oxidized and reduced glutathione levels in Mn-depleted *S. sanguinis* cells

Glutathione (γ-glutamyl-cysteinylglycine) is a nonprotein thiol produced by cells to prevent damage caused by reactive oxygen species (ROS) (Carmel-Harel and Storz, 2000; Sies, 1999). The SK36 genome (Xu *et al*., 2007) encodes a bifunctional γ-glutamate-cysteine ligase/glutathione synthetase (GshF; SSA_2168) (Janowiak and Griffith, 2005). Mean levels of the glutathione precursors cysteine, glutamine, and γ-glutamylcysteine all decreased slightly in cells after Mn depletion, consistent with active synthesis, although glycine levels did not change **(Figure 5a)**. Interestingly, levels of reduced glutathione (GSH) increased in cells after Mn depletion, whereas levels of the oxidized form (GSSG) remained constant **(Figure 5b)**. Since the air flow was kept constant throughout the experiment, we expected that GSH would have been utilized by redox enzymes for ROS remediation. While ROS levels were not measured directly by the metabolomics analysis, levels of *ortho*-tyrosine increased (**Figure 5c**), which is an indicator of high ROS states (Ipson and Fisher, 2016; Matayatsuk *et al*., 2007). Thus, the accumulation of GSH is probably due to Mn depletion, either because of a blockage of GSH utilization by redox enzymes or due to a reduction of ROS.

**Fig. 5.**
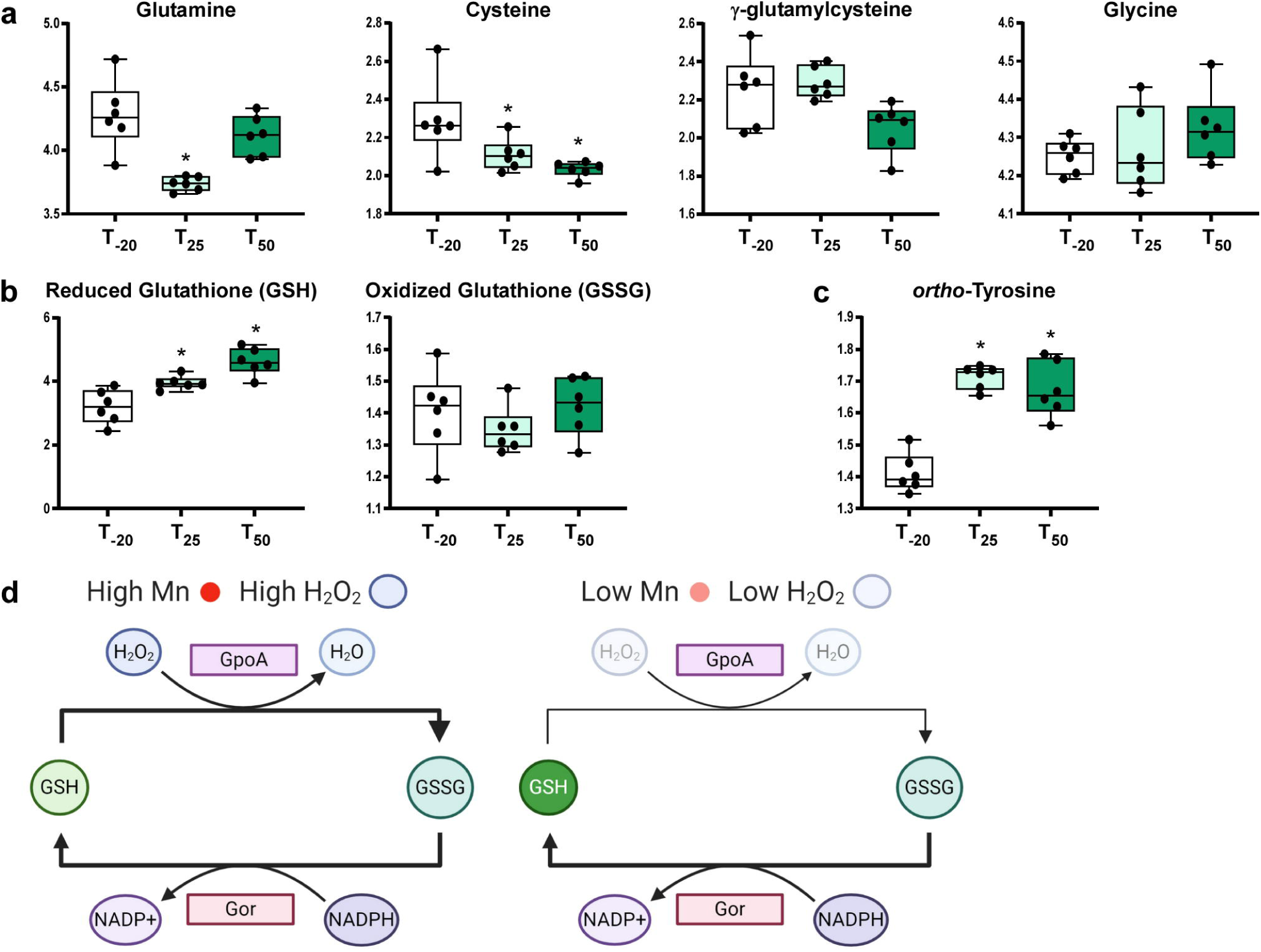
Glutathione abundance in cells and model of Mn depletion. Levels of glutathione precursors (a), glutathione (b), and the oxidative stress indicator *ortho-*tyrosine (c) are shown. Metabolite levels were compared using one-way ANOVA with a Fisher’s least significant difference test to compare the post-EDTA samples to pre-EDTA. Black asterisks indicate P-value < 0.05. (d) Model of glutathione utilization by glutathione peroxidase (GpoA) and reduction by glutathione reductase (Gor) under normal and low-Mn conditions. Figure (d) was made using Biorender.com.

Due to the presumed decrease in activity of the Mn-cofactored superoxide dismutase, SodA (Crump *et al*., 2014), it is unlikely that all ROS levels would have decreased after Mn depletion. The notable exception is H_2_O_2_, which was found to decrease after Mn depletion due to reduced expression of *spxB* (Puccio *et al*., 2020). This likely led to a decrease in the direct detoxification of H_2_O_2_ by GSH, although the extent to which this occurs in cells is controversial (Berndt *et al*., 2014). Additionally, *S. sanguinis* does not encode any known glutaredoxins and the only enzyme thought to utilize GSH in *S. sanguinis* is glutathione peroxidase (GpoA; SSA_1523), which uses GSH to detoxify H_2_O_2_ (**Figure 5d**) (Carmel-Harel and Storz, 2000). This enzyme has been found to contribute to oxidative stress tolerance in *S. pneumoniae* (Potter *et al*., 2012) and virulence in *S. pyogenes* (Brenot *et al*., 2004). Additionally, the enzyme that converts GSSG to GSH, glutathione reductase (Gor; SSA_1533), is likely metal-cofactored, which could explain why GSSG levels remained constant instead of decreasing as GSH levels increased. Thus, Mn depletion could explain the accumulation of both reduced and oxidized glutathione.

## 5. Conclusions

In this study, we showed system-wide metabolomic changes induced in *S. sanguinis* Mn-transporter mutant cells and spent media in response to EDTA treatment over time. This study captured the Mn-responsive metabolic processes, such as dysregulations in carbohydrate, nucleotide, and redox metabolism, many of which may contribute to the reduction in bacterial growth rate and virulence. The decrease in available Mn led to the accumulation of fructose-1,6-bisphosphate, which likely resulted in induction of carbon catabolite repression. This has widespread consequences, such as the blockage of nucleobases conversion into nucleosides and accumulation of reduced glutathione. In addition, we provide insights into the metabolic composition of BHI and the components streptococci may utilize from this undefined medium.

## 6. Declarations

### Ethics approval

This article does not contain any studies with human participants or animals performed by any of the authors.

### Consent to participate

Not applicable

### Consent for publication

All authors have read, approved and have provided consent for this publication.

### Availability of data and material

The datasets generated and analyzed during the current study are available as Supplementary Tables S1 and S2 as provided by Metabolon, Inc.

## Code availability

Not applicable

## Author’s contributions

TP and TK designed the experiments. TP performed the experiments. BBM performed the data analysis. All authors analyzed the results and wrote the manuscript.

## Acknowledgements

We thank Karina Kunka, Dr. Shannon Green, Dr. Seon-Sook An, and Brittany Spivey for discussions and assistance with experiments. We also thank Dr. Danny Alexander (Metabolon, Inc.) for his initial analysis of the data.

## Supplementary Table Captions

**Supplementary Table S1**. Raw metabolite abundance data for cellular metabolites captured using combined LC-MS/MS (positive and negative modes) analysis by Metabolon, Inc. Retention indices (RIs), quantifier mass, CAS IDs, KEGG IDs, HMDB IDs, PubChem IDs, SMILES, Super Pathway and Sub Pathway information, biochemical names for the metabolites, and their raw abundances are also included.

**Supplementary Table S2**. Raw metabolite abundance data for media metabolites captured using combined LC-MS/MS (positive and negative mode) analysis by Metabolon, Inc. Retention indices (RIs), quantifier mass, CAS IDs, KEGG IDs, HMDB IDs, PubChem IDs, SMILES, Super Pathway and Sub Pathway information, biochemical names for the metabolites, and their raw abundances are also included.

**Supplementary Table S3**. Transformed, scaled and normalized metabolite abundance data for cellular metabolites.

**Supplementary Table S4**. Transformed, scaled and normalized metabolite abundance data for media metabolites.

**Supplementary Table S5**. Pathway enrichment analysis for the 534 quantified cellular metabolites.

**Supplementary Table S6**. Pathway enrichment analysis for the 424 quantified metabolites in media.

**Supplementary Table S7**. Unique metabolites found in some but not all samples.

**Supplementary Table S8**. One-way ANOVA statistical analysis results for metabolites of cells. Fold changes cut off > 1.2 and < 0.8; P-value < 0.05.

**Supplementary Table S9**. One-way ANOVA statistical analysis results for metabolites of media. Fold changes cut off > 1.2 and < 0.8; P-value < 0.05).

**Supplementary Table S10**. Pathway enrichment analysis for significantly differential (ANOVA) cellular metabolites.

**Supplementary Table S11**. Pathway enrichment analysis for significantly differential (ANOVA) media metabolites.

**Supplementary Table S12**. Average cell metabolite levels, fold changes, and P-values as determined by t-tests comparing post-EDTA samples to the pre-EDTA sample.

**Supplementary Table S13**. Average media metabolite levels, fold changes, and P-values as determined by t-tests comparing spent media (T_-20_) to pre-inoculation media as well as post-EDTA samples to the pre-EDTA sample.

**Supplementary Table S14**. STEM analysis of cellular metabolites displaying top 2 significant profiles.

**Supplementary Table S15**. STEM analysis of media metabolites displaying top 3 significant profiles.

