## Supplementary Material for "Time-course analysis of *Streptococcus sanguinis* after manganese depletion reveals changes in glycolytic, nucleotide, and redox metabolites"

Tanya Puccio[1], Biswapriya B. Misra[2], Todd Kitten[1]*

[1]Philips Institute for Oral Health Research, Virginia Commonwealth University School of Dentistry, Richmond 23298, VA USA.
[2]Department of Internal Medicine, Section on Molecular Medicine, Wake Forest School of Medicine, Medical Center Boulevard, Winston-Salem 27157, NC USA.

***Correspondence:**
Dr. Todd Kitten

804-628-7010

### Supplementary Methods

#### Sample preparation

Samples stored at -80°C, upon shipment were accessioned into the Metabolon LIMS system were prepared using the automated MicroLab STAR® system from Hamilton Company. Several recovery standards were added prior to the first step in the extraction process for QC purposes. Samples were extracted with methanol under vigorous shaking for 2 min (Glen Mills GenoGrinder 2000) to precipitate protein and dissociate small molecules bound to protein or trapped in the precipitated protein matrix, followed by centrifugation to recover chemically diverse metabolites. The resulting extract was divided into five fractions: two for analysis by two separate reverse phase (RP)/UPLC-MS/MS methods using positive ion mode electrospray ionization (ESI), one for analysis by RP/UPLC-MS/MS using negative ion mode ESI, one for analysis by HILIC/UPLC-MS/MS using negative ion mode ESI, and one reserved for backup. Samples were placed briefly on a TurboVap® (Zymark) to remove the organic solvent. The sample extracts were stored overnight under nitrogen before preparation for analysis.

#### Metabolomics data generation using Ultrahigh Performance Liquid Chromatography-Tandem Mass Spectroscopy (UPLC-MS/MS)

All methods utilized a Waters ACQUITY ultra-performance liquid chromatography (UPLC) and a Thermo Scientific Q-Exactive high resolution/accurate mass spectrometer interfaced with a heated electrospray ionization (HESI-II) source and Orbitrap mass analyzer operated at 35,000 mass resolution. The sample extract was dried then reconstituted in solvents compatible to each of the four methods. Each reconstitution solvent contained a series of standards at fixed concentrations to ensure injection and chromatographic consistency. One aliquot was analyzed using acidic positive ion conditions, chromatographically optimized for more hydrophilic compounds. In this method, the extract is gradient-eluted from a C18 column (Waters UPLC BEH C18-2.1x100 mm, 1.7 µm) using water and methanol, containing 0.05% perfluoropentanoic acid (PFPA) and 0.1% formic acid (FA). A second aliquot was also analyzed using acidic positive ion conditions, but chromatographically optimized for more hydrophobic compounds. In this method, the extract is gradient eluted from the aforementioned C18 column using methanol, acetonitrile, water, 0.05% PFPA and 0.01% FA, and is operated at an overall higher organic content. A third aliquot was analyzed using basic negative ion optimized conditions using a separate dedicated C18 column. The basic extracts were gradient-eluted from the column using methanol and water, however with 6.5mM Ammonium Bicarbonate at pH 8. The fourth aliquot was analyzed via negative ionization following elution from a HILIC column (Waters UPLC BEH Amide 2.1x1$_{50}$ mm, 1.7

µm) using a gradient consisting of water and acetonitrile with 10 mM Ammonium Formate, pH 10.8. The MS analysis alternated between MS and data-dependent $MS^n$ scans using dynamic exclusion. The scan range varies slightly between methods, but covers approximately 70-1000 m/z. Raw data files were archived and extracted as described below.

**Data Extraction and Compound Identification**

Raw data were extracted, peak-identified, and QC processed using Metabolon's hardware and software. These systems are built on a web-service platform utilizing Microsoft's .NET technologies, which run on high-performance application servers and fiber-channel storage arrays in clusters to provide active failover and load-balancing. Compounds were identified by comparison to library entries of purified standards or recurrent unknown entities. Metabolon maintains a library based on authenticated standards that contains the retention time/index (RI), mass to charge ratio (*m/z)*, and chromatographic data (including MS/MS spectral data) on all molecules present in the library. Furthermore, biochemical identifications are based on three criteria: retention index within a narrow RI window of the proposed identification, accurate mass match to the library +/- 10 ppm, and the MS/MS forward and reverse scores. MS/MS scores are based on a comparison of the ions present in the experimental spectrum to ions present in the library entry spectrum. While there may be similarities between these molecules based on one of these factors, the use of all three data points can be utilized to distinguish and differentiate biochemicals. More than 4500 commercially available purified standard compounds have been acquired and registered into LIMS for analysis on all platforms for determination of their analytical characteristics. Additional mass spectral entries have been created for structurally unnamed biochemicals, which have been identified by virtue of their recurrent nature (both chromatographic and mass spectral). Putative identification of each metabolite was made based on mass accuracy (m/z) Chemical Abstracts Service (CAS), Kyoto Encyclopedia of Genes and Genomes (KEGG), Human Metabolome Database (HMDB), and LIPID MAPS identifiers.

**Curation**

A variety of curation procedures were performed to ensure that a high quality data set was made available for statistical analysis and data interpretation. The QC and curation processes were designed to ensure accurate and consistent identification of true chemical entities, and to remove those representing system artifacts, mis-assignments, redundancy, and background noise. Metabolon data analysts used internally-developed visualization and interpretation software to confirm the consistency of peak identification among the various samples. Library matches for each compound were checked for each sample and corrected if necessary.

### Supplementary Figures

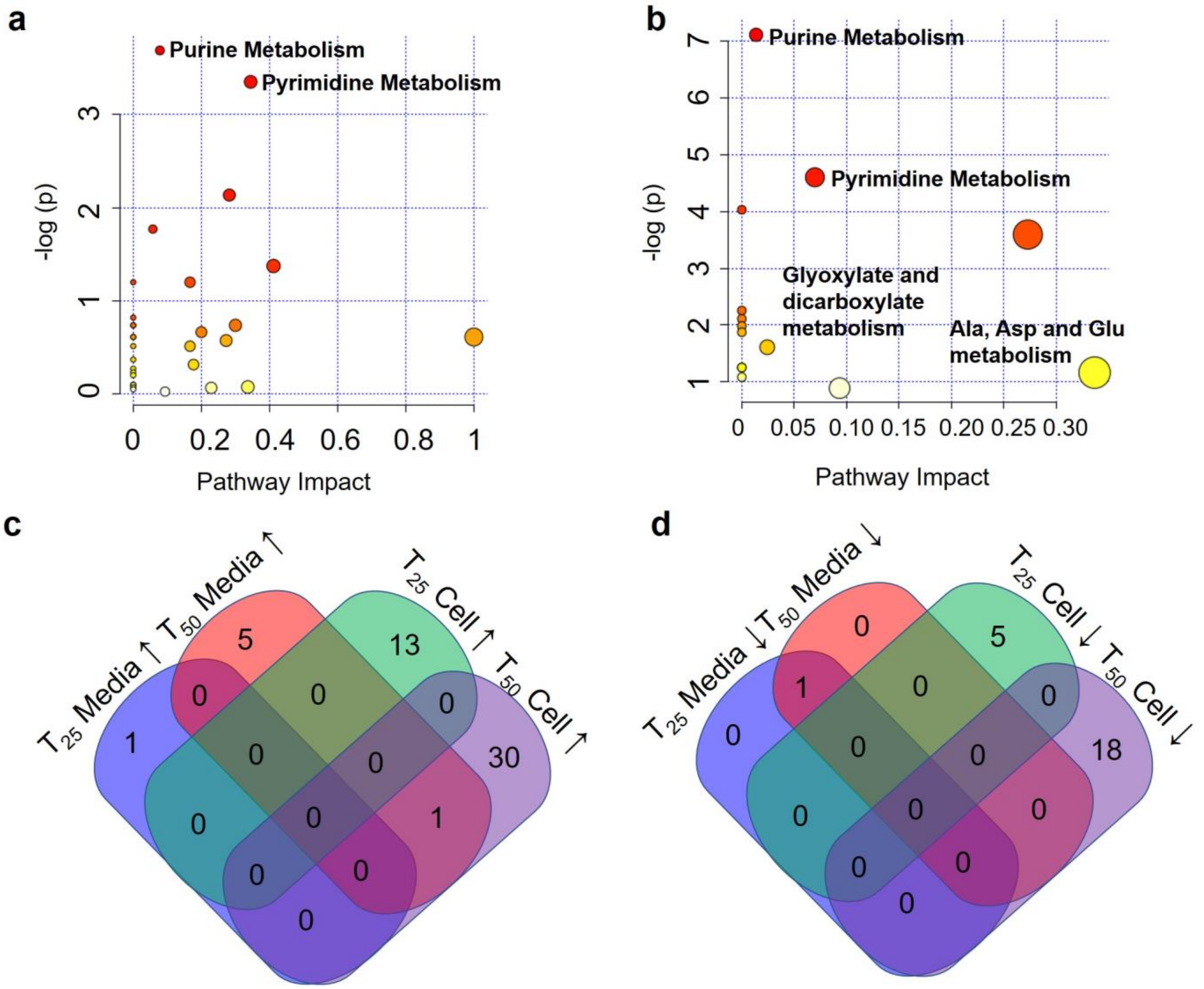


**Figure S1. Pathway enrichment analysis for differential metabolites**

Pathway enrichment analysis for significantly differential metabolites (ANOVA) in cells (a) and spent media (b). A 4-way Venn diagram displaying significantly increased (c) and decreased (d) metabolites at $T_{25}$ and $T_{50}$ compared to $T_{-20}$.

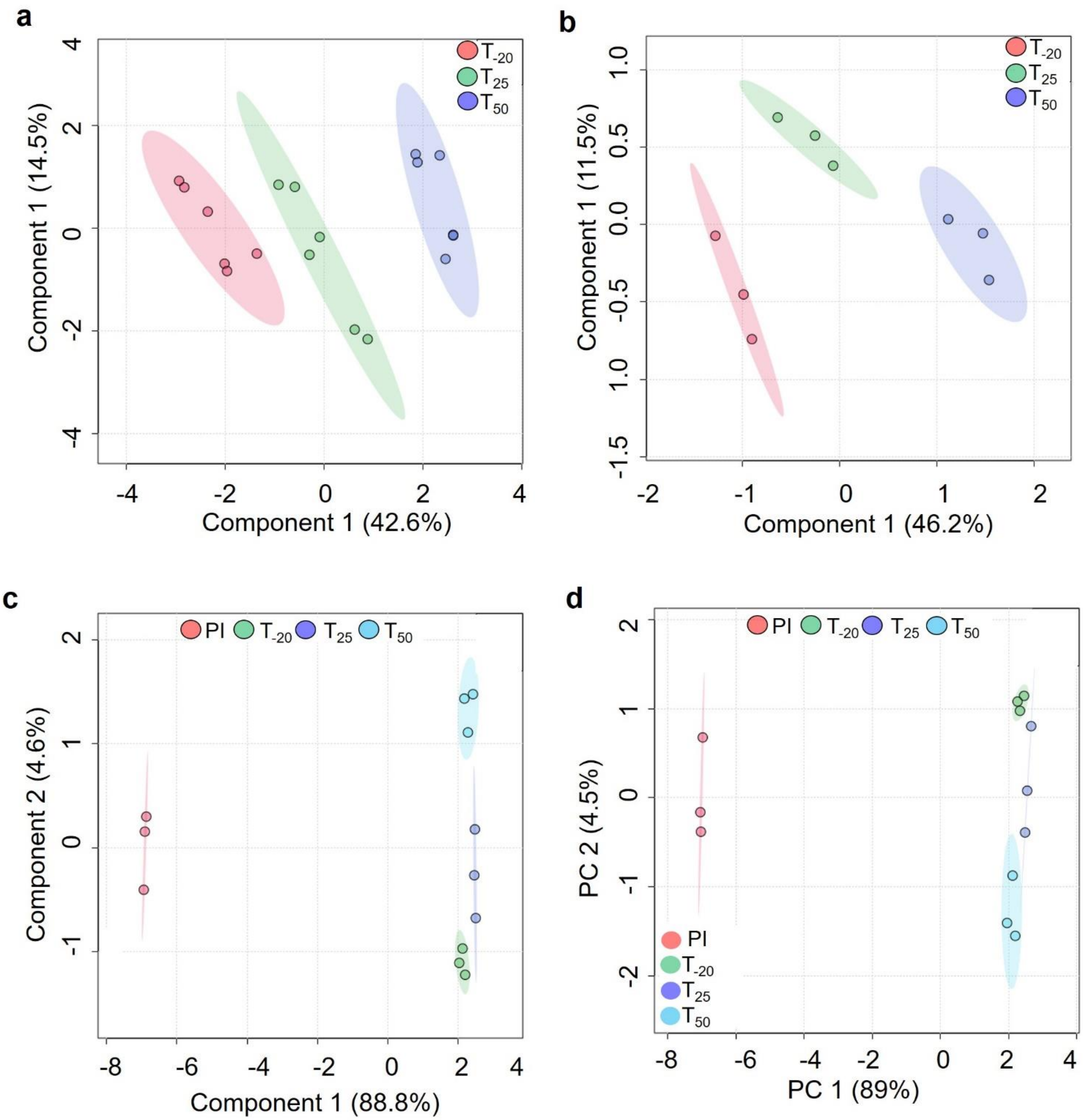


**Figure S2. Multivariate (PLS-DA) analysis of the metabolomic changes in cells and media.**

(a) Score plot for PLS-DA displaying the separation of time-points in cells (a) and spent media (b). Multivariate analysis of all media samples using PLS-DA (c) and PCA (d).

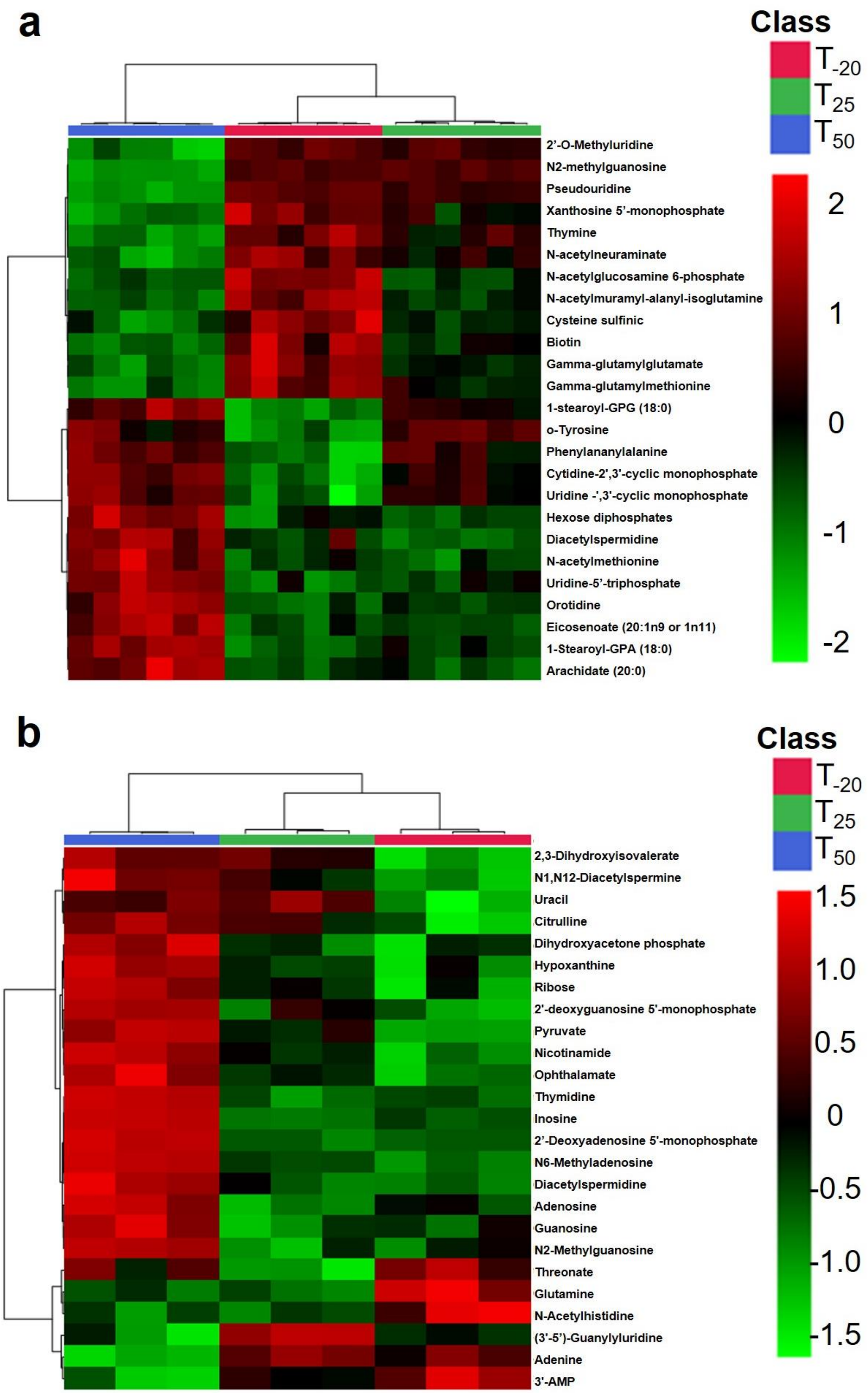


**Figure S3. Hierarchical clustering analysis of the metabolites of cells and media.**

Top 25 ANOVA-derived differential metabolites for HCA in cells (a) and spent media (b).

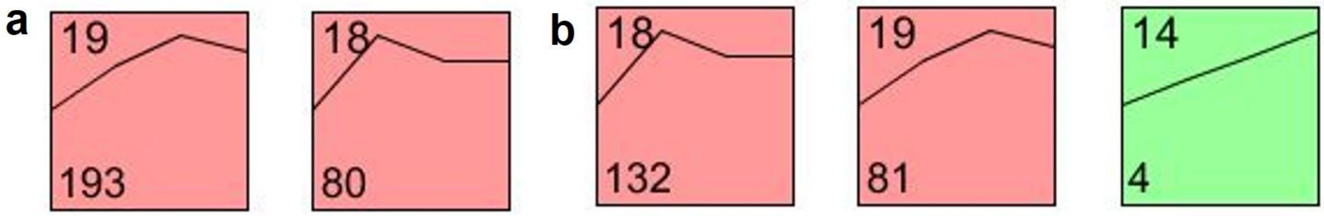


**Figure S4. Time course analysis of cellular and media metabolism.** Models displaying the time-dependent changes in metabolite abundance in cells (a) and spent media (b). Models #19 and #18 were statistically significant in cells among the 20 models interrogated. Models # 18, #19, and #14 were statistically significant in media.

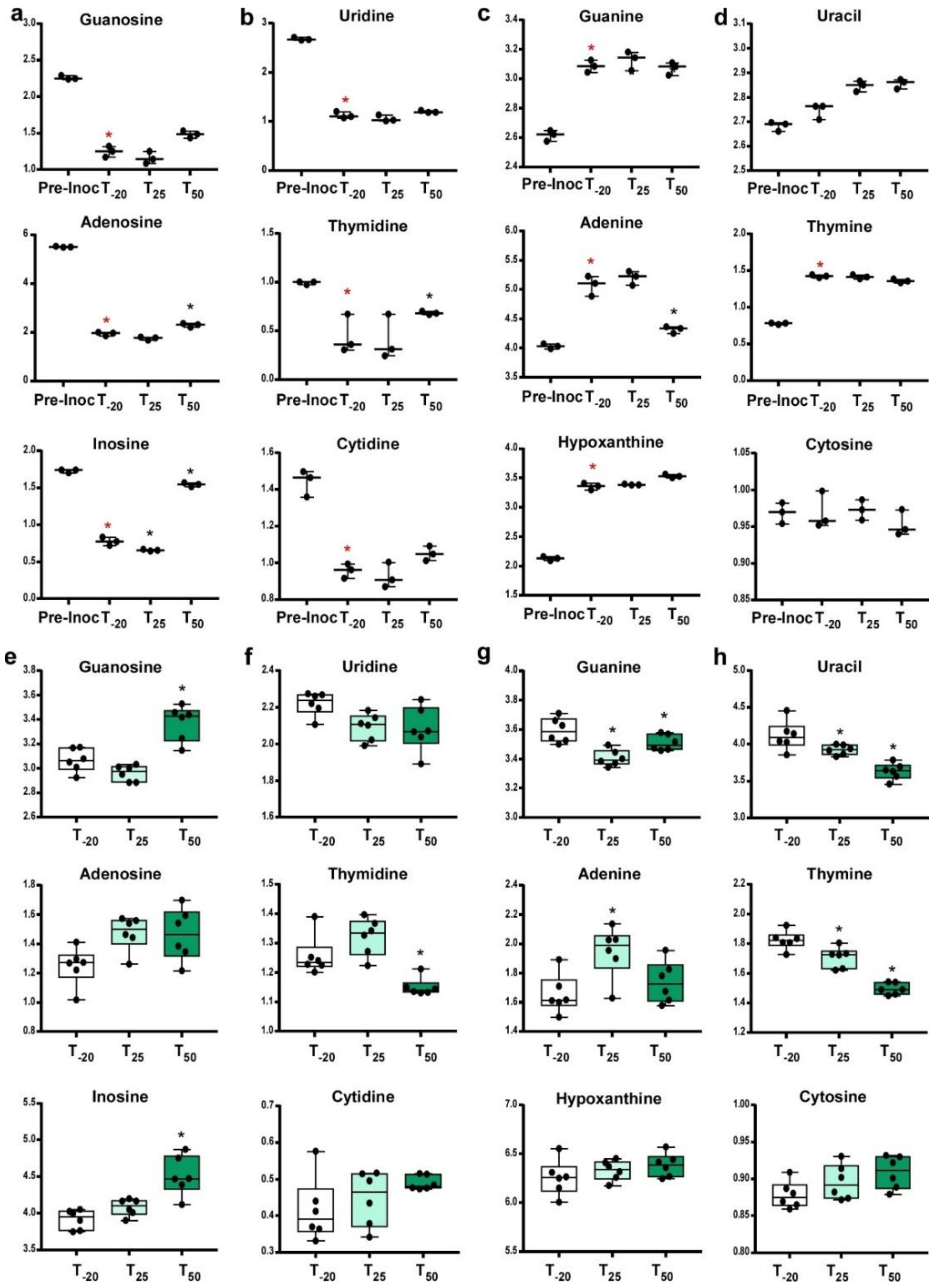


**Figure S5 Quantitative changes in nucleotide metabolism**

Purine nucleosides in media (a) and cells (e). Pyrimidine nucleosides in media (b) and cells (f). Purine nucleobases in media (c) and cells (g). Pyrimidine nucleobases in media (d) and cells (h). Whiskers indicate the range; horizontal bars represent the mean. A two-tailed t-test was used to compare the pre-inoculum (Pre-Inoc) media samples to post-inoculum ($T_{-20}$). Red asterisks indicate when P-value < 0.05. Spent media and cell metabolite levels were compared using an ANOVA with a Fisher's least significant difference test to compare the post-EDTA samples to pre-EDTA. Black asterisks indicate P-value < 0.05.
